# Targeted knockdown of the PAF49 component of the PAF53/PAF49 heterodimer causes the degradation of PAF53

**DOI:** 10.1101/2021.01.27.428437

**Authors:** Rachel McNamar, Katrina Rothblum, Lawrence I. Rothblum

**Author notes:** To whom correspondence should be addressed: Lawrence Rothblum, Department of Cell Biology, BMSB 553, University of Oklahoma College of Medicine, Oklahoma City, OK. 73104.

## Abstract

There are significant differences in the components of the ribosomal DNA transcription apparatuses of yeast and mammals. Moreover, the patterns of regulation between mammals and yeast are also different. To overcome, deficits in our understanding of mammalian rDNA transcription, we have developed a system to introduce an inducible degron into the endogenous genes of mammalian cells. This allows us to combine a knock out the endogenous gene product and replace it with mutant proteins in order to study their function in ribosomal DNA transcription. Using this system, we show that the knockout of PAF49, the mammalian ortholog of yeast A34, results in the relatively rapid degradation of PAF53, the ortholog of yeast A49. Interestingly, the steady-state levels of the core subunits of RNA polymerase I are unaffected.

---

When deprived of nutrients and/or growth factors, mammalian cells repress rDNA transcription by as much as 80–90%. Similarly, when nontransformed cells are grown to confluence, rDNA transcription is also turned down. The downshift in rDNA transcription can be accomplished through various mechanisms or signaling pathways. These mechanisms can include changes in the post-translational modification or changes in the steady-state levels of the components of the transcription apparatus. Importantly, the mechanisms used to turn down rRNA synthesis can vary between different cell types (1-14) (2,12,15).

Our studies focus on the complex regulation of two additional Pol I components. These two Pol I subunits, A34.5p (A34, PolR1G) and A49p (A49, PolR1E) form a heterodimer that binds to the lobe of the polymerase structure and can be dissociated from the core polymerase. The proteins have poorly defined roles in rDNA transcription. *S. cerevisiae* A49 is reported to not be *essential* for viability (16) or rDNA transcription. RPA49 Δ yeast grow at 6% of the wild type rate at 25C (16). Similarly, deletion of the *S. pombe* homologue, RPA51 (17), reduced rDNA transcription 70% (no effect on nonspecific polymerase activity) and reduced the rate of growth 50% at 30C.

Deletion of the other partner in the heterodimer, yA34, has a minor effect on growth and/or rRNA synthesis. However, deletion of A34 appears to weaken the binding of A49 to the core Pol I, as polymerase loses the A49 subunit upon purification (18,19). Biochemical purification of wild type yeast Pol I results in two fractions, one of which does not contain either A49 or A34 (20). This is complemented by the observation that a large fraction of the polymerase particles prepared for cryo-EM “either lacked the A49/A34.5 subcomplex or displayed a flexible clamp-stalk region” (21). Interestingly, cross-linking experiments indicate that most of the interactions between the heterodimer and yeast Pol I appear to be mediated by the A49 subunit (22). Portions of the N-termini of A34 and A49 form a heterodimerization domain that stimulates Pol I RNA cleavage and processivity. The heterodimerization domain forms a fold with structural similarity to the core of the Pol II general transcription factor TFIIF (23), which similarly binds to the lobe of the polymerase. The C-terminus of yeast A49 contains a tandem winged helix domain (tWH) (23) that is capable of DNA-binding and resembles a similar element in TFIIE and the Pol III TFIIE-like Rpc34.

Mammalian cells contain orthologs to yeast A49 and A34. PAF53 and PAF49, the orthologs of A49 and A34, respectively, were first identified in Muramatsu’s laboratory. That laboratory reported that PAF49 and PAF53 are associated with a fraction of the core Pol I molecules and were essential for promoter-specific transcription (24,25). This observation was confirmed and extended by Hannan *et al*. who estimated that only 60% of the polymerase molecules in a rat hepatoma cell line contained PAF53 (26). Muramatsu’s laboratory reported that the nucleolar localization of PAF49 was growth– dependent. This result was subsequently confirmed by our laboratory (27-29). We reported that the nucleolar levels of both PAF53 and PAF49 decreased when cells were serum starved. More recently, PAF53 and PAF49 were identified as “being essential” in CRISPR/Cas9 screenings of the mammalian genome (30,31). This was confirmed in a directed CRISPR experiment when we found that following attempts to eliminate PAF53 expression using CRISPR, the only clones we could isolate expressed PAF53 (32), but also contained evidence of a recombined gene.

The Pol I-specific “**p**olymerase **a**ssociated **f**actors” (Rrn3 and the heterodimer of PAF49 and PAF53) assemble with Pol I in the prelude to rDNA transcription (3-5,9-15,17,19,24,25,28,33-48). While KO studies in mammalian cells demonstrate an essential role for the PAF53/PAF49 complex (heterodimer) in rDNA transcription and cellular physiology (23,30,32,49), their roles in transcription are unknown. Moreover, there are different patterns of regulation between mammals and yeast. Thus, although studies on yeast rDNA transcription can serve as a model for mammalian rDNA transcription, there are significant deficits in our understanding of mammalian rDNA transcription. To facilitate the study of rDNA transcription in mammalian cells, we developed a system using an auxin-inducible degron that allows us to specifically and rapidly knockdown the levels of a target protein and replace it with mutants. This system enables us to combine a “genetics-like” approach to studying mammalian rDNA transcription with biochemistry. Using this system, we have found that PAF53 is required for rDNA transcription and mitotic growth and we were able to establish that the dimerization, linker and tWH domains of PAF53 are involved in rDNA transcription. Further, both the linker and tWH helix domains of mammalian PAF53 were found to have DNA-binding activity.

We have previously reported that cells appeared to co-regulate PAF53 and PAF49 levels. For example, when 3T6 cells were serum starved, we observed a 70% decrease in the levels of both proteins. Similarly, when 3T3 cells were serum starved, we observed a 30% decrease in both proteins (29). Interestingly, studies on yeast A34 and A49 have not reported the coordinated regulation of the steady-state levels of these proteins except to note that interactions between the two affected their interaction with Pol I (19). On the other hand, it has been reported that the interaction of yeast A49 with Pol I is apparently independent of its interaction with A34 as it does not require the dimerization domain (23,50).

In the course of our investigation of the roles of PAF53 and PAF49 in rDNA transcription using an auxin-induced degron, we considered the possibility that the induced knockdown of one component of the heterodimer might affect the cellular level of its partner. We now report that the induced degradation of PAF49 caused a decrease in PAF53, but the reverse did not occur. Degradation of the targeted protein is through the auxin induced activation of a plant E3 ligase (TIR1) and should result in proteasomal-dependent degradation of the targeted protein. Interestingly, the proteasome inhibitor MG132 inhibited the AID-tagged PAF49 and also inhibited the degradation of PAF53. Genetic studies in yeast have not observed a linkage in the expression of the two partners.

Our results suggest that in mammalian cells, PAF49 and PAF53 are coregulated post-translationally and that the regulation of their stability in the cells would represent a new mechanism for regulating rDNA transcription in mammals

## Results

Previously, we targeted PAF53 for degradation and replaced the protein with mutants so that we could map functional domains. In yeast, the knockout of one of the components of the A49/A34 heterodimer has not been reported to affect the expression of the other (references). However, based on the yeast model, we did not determine if the loss of PAF53 would destabilize PAF49. Hence, we sought to confirm the observations made in yeast in mammalian cells.

We targeted PAF53 for degradation as described previously (51) and determined the level of PAF49. As shown in Figure 1A, the knockdown of PAF53 for 24 hours is not associated with the depletion of PAF49 (compare lanes 1 and 2). We then carried out the reciprocal experiment, targeting PAF49 for auxin-dependent degradation. We found that the knockdown of PAF49 caused a rapid attenuation of PAF53 levels in several independent clones expressing AID-PAF49. To further investigate this phenomenon, we examined the effects of knocking down either PAF53 or PAF49 on the stability of the core Pol I subunits.

**Figure 1.**
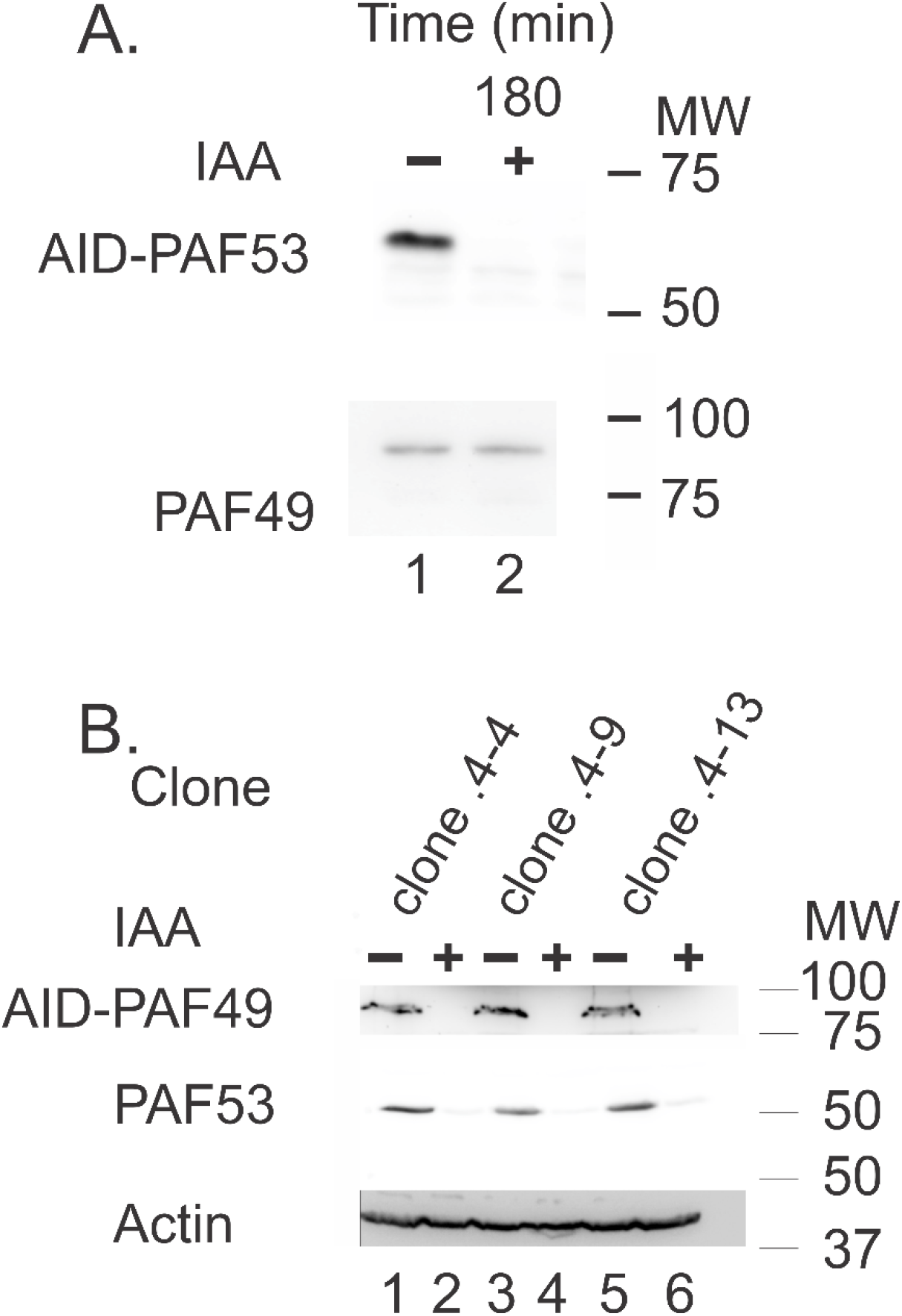
Indole acetic acid (IAA) induces the degradation of AID-tagged PAF53 (A) and AID-tagged PAF49 (B) in cell lines in which the genes have been tagged to express AID-tagged proteins. (B) The IAA induced degradation of AID-PAF49 results in the degradation of PAF53 in the same cells. The analyses presented were carried out after three hours of treatment with 500 μM IAA (+) or water (−). Cells were harvested and western blots for PAF53 and PAF49 were carried out as described (26,51).

The knockdown of PAF53 had no significant effect on the steady-state levels of either A194, A127 or rpa43 (Figure 2A). Similarly, the induced degradation of PAF49 had no effect on the levels of the other polymerase subunits (Figure 2B). Interestingly, while the induced degradation of PAF53 did not have an immediate effect on the level of PAF49, we did observe a significant decrease in PAF49 levels after 72 hours (Figure 3). RT-qPCR analysis of the levels of PAF53 and PAF49 mRNA did not demonstrate a significant change in their levels after 24 hr.

**Figure 2.**
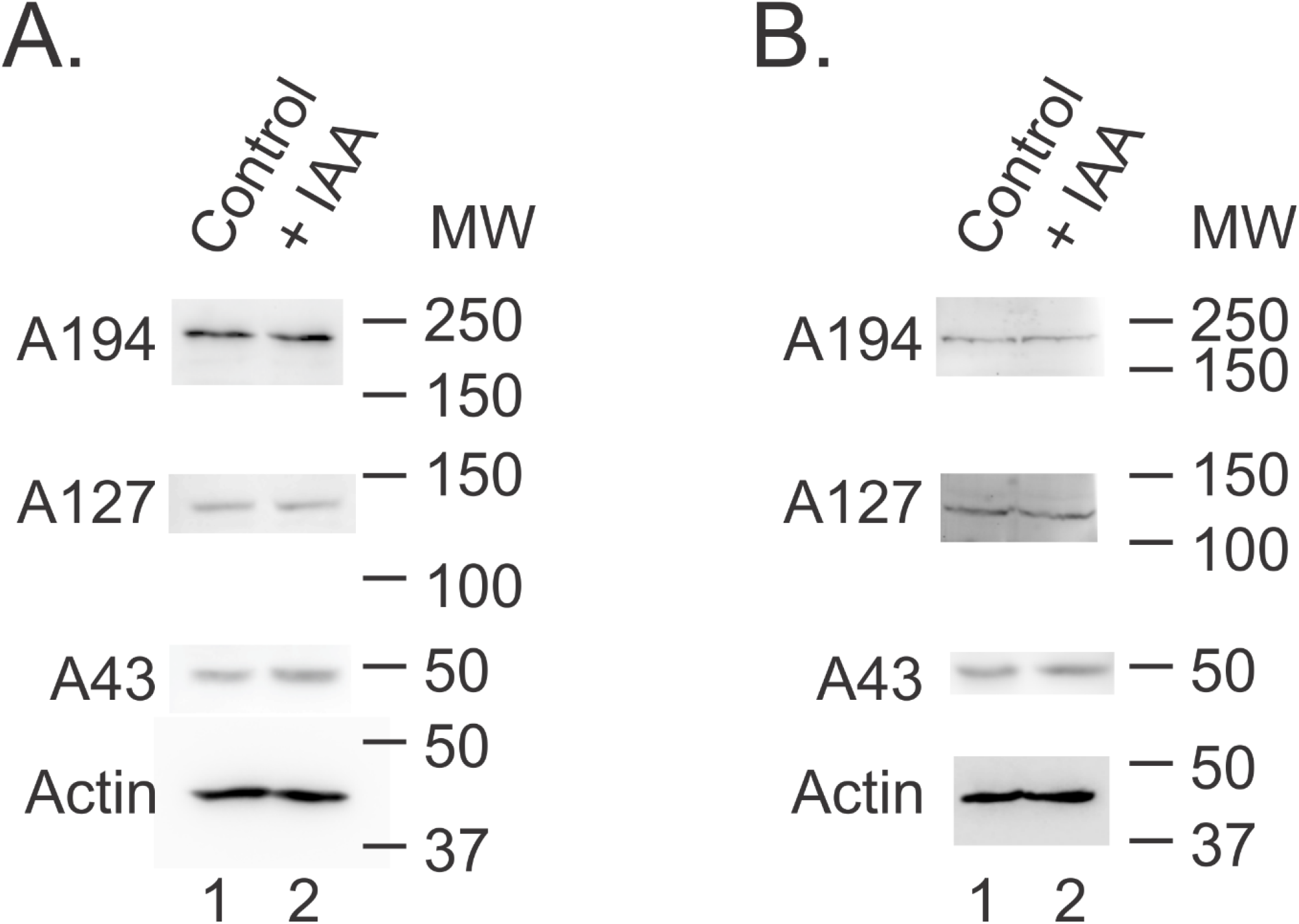
After three hours, the knockdown of either AID-PAF53 (A) or AID-PAF49 (B) has no effect on the stability of core Pol I subunits. Cells expressing AID-tagged PAF53 (A) or AID-tagged PAF49 (B) were treated with 500 μM IAA (+) or water (-) for the indicated lengths of time, after which they were harvested and analyzed by western blotting as described in Materials and Methods. Experimental procedures are as described in the legend to Figure 1.

**Figure 3.**
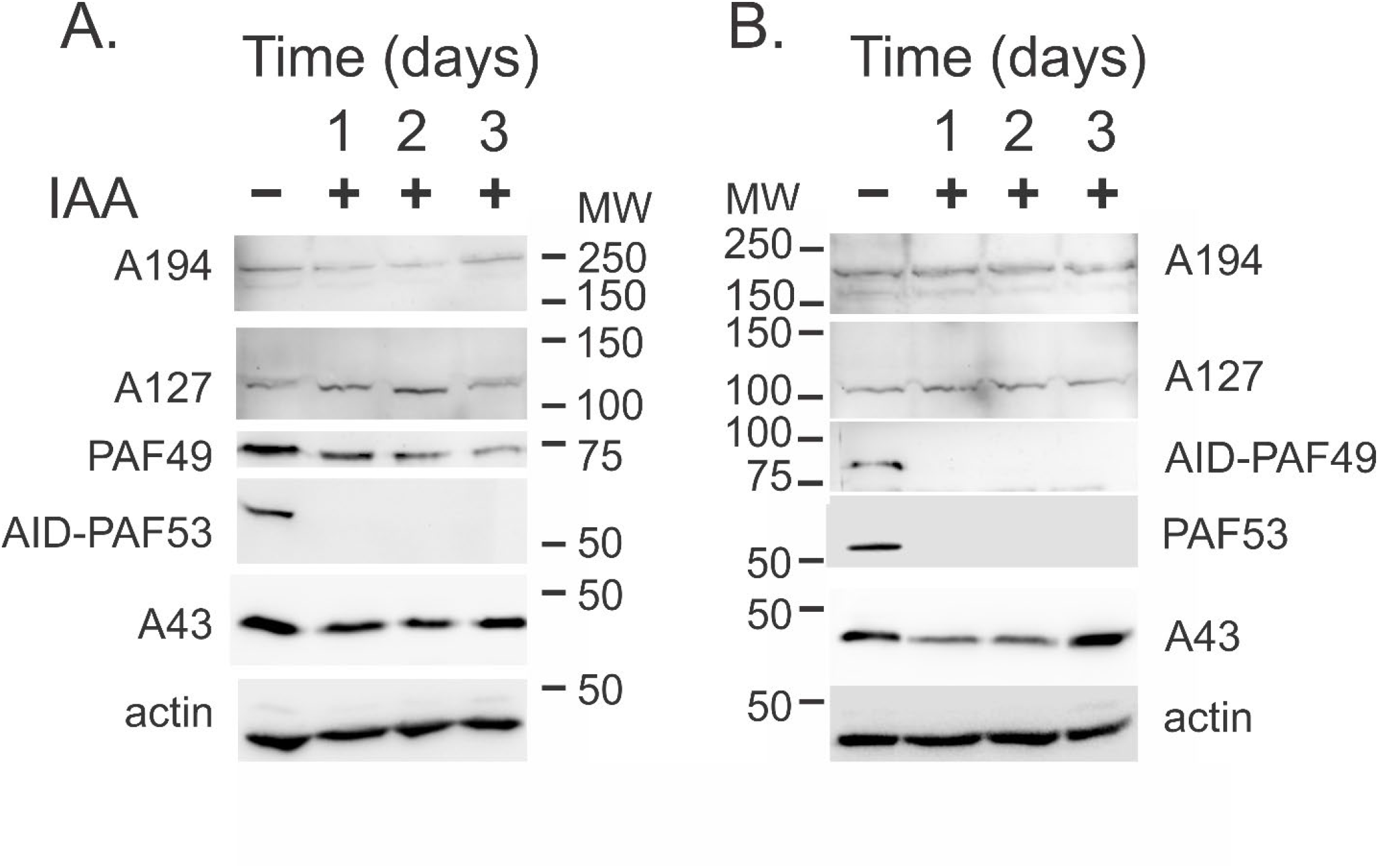
The prolonged knockdown of AID-PAF53 (A) or AID-PAF49 (B) has no effect on the steady-state levels of the core Pol I subunits, A194 (POLR1A), A127 (POLR1B) and A43 (RPA43, POLR1F). Cells expressing AID-tagged PAF53 (A) or AID-tagged PAF49 (B) were treated with 500 μM IAA (+) or water (-) for the indicated lengths of time, after which they were harvested and analyzed by western blotting as described in Materials and Methods.

We next compared the time courses of the degradation of the two proteins. As shown in Figure 4A (repeated from Figure 1), the rapid degradation of PAF53-AID had little effect on the steady state level of PAF49 after 180 minutes. In contrast, rapid degradation of AID-PAF49 was paralleled by the disappearance of PAF53 (Figure 4B). Figure 4C summarizes the kinetics of the changes in the levels of the two proteins as seen in three independent experiments.

**Figure 4.**
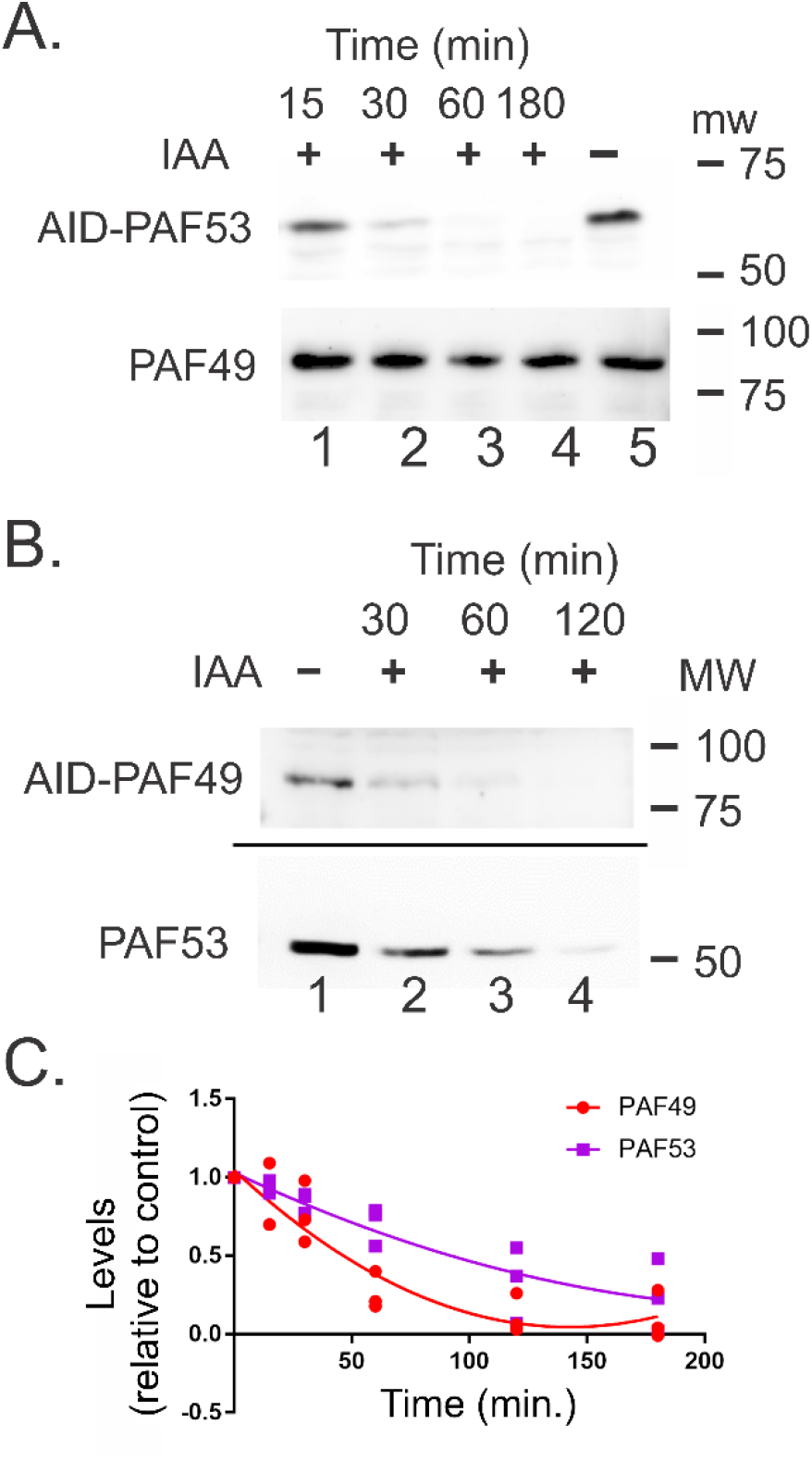
The IAA-induced degradation of AID-PAF49 results in the knockdown of PAF53 in the same cells. (A) The IAA-induced degradation of AID-PAF53 does not cause a reduction in PAF49. (B) The IAA-induced degradation of AID-PAF49 results in the rapid, time-dependent knockdown in PAF53. Cells were treated with 500 μM IAA (+) or water (-) for the indicated length of time, and then harvested and analyzed by western blotting as indicated (26,51). (C) Quantitation of the cellular levels of AID-PAF49 and PAF53 following treatment with IAA (n=3).

One explanation for the parallel degradation of the components of the heterodimer would be that they are both being degraded through the same pathway. The auxin inducible degron that we are using in these experiments is recognized by TIR1, an E3 ubiquitin ligase that is auxin responsive. This should result in the ubiquitin-dependent, proteasomal degradation of the target proteins. Hence, the knockdown of the AID-tagged proteins should be inhibited by agents that inhibit the proteasome such as MG132 (52). When 293 cells expressing AID-PAF53 were treated with IAA and MG132, the degradation of PAF53 was inhibited. (Figure 5A). When we carried out the reciprocal experiment, we observed a parallel result. The auxin-dependent degradation of AID-PAF49 was inhibited by treatment with MG132. Interestingly the decrease in nontagged-PAF53 was also inhibited by MG132 (Figure 5B).

**Figure 5.**
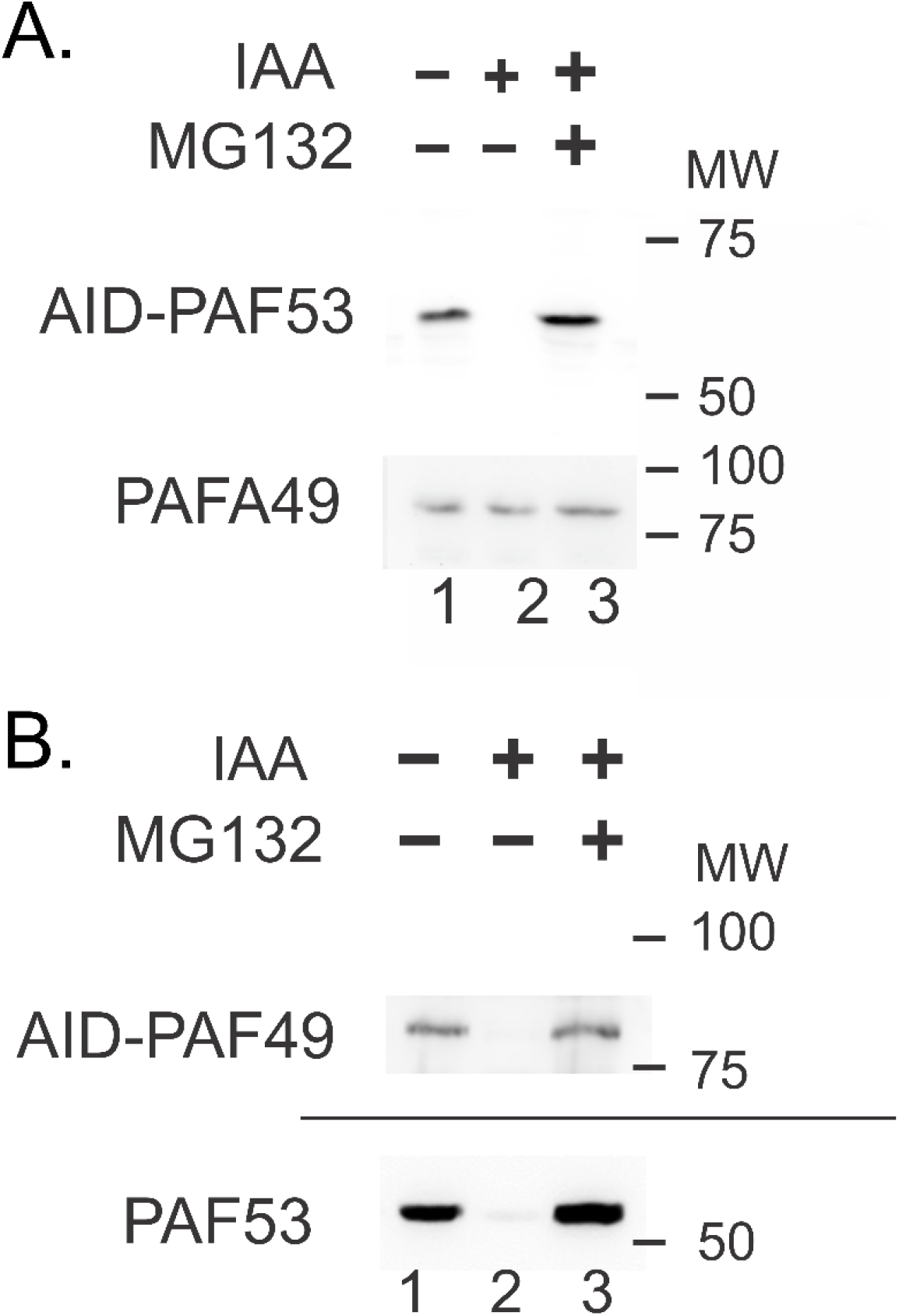
MG132 blocks the IAA-induced degradation of AID-PAF53 (A) and AID-PAF49 (B). (B) MG132 blocks the IAA induced degradation of untagged PAF53 in cells expressing AID-PAF49. Experimental procedures are as described in the legend to Figure 1. When indicated, the MG132 (10 μM) was added to the cell cultures at the same time as IAA (500 μM).

These data suggested that the programmed, proteosomal-dependent degradation of PAF49 of the heterodimer induced the degradation of PAF53 However, it does not distinguish between possible mechanisms of degradation of the partner, *i*.*e*. a ubiquitin-dependent pathway of degradation or a non-ubiquitin-dependent pathway.

## Discussion

Muramatsu’s laboratory was the first to report the coordinated regulation of PAF53 and PAF49. They found that serum starvation caused the loss of both proteins from the nucleoli of NIH 3T3 cells (24). Our laboratory confirmed these findings and reported that both the nucleolar localization and steady-state levels of the two proteins were coregulated (27,29). We found that when NIH3T6 cells were serum starved, the levels of both PAF53 and PAF49 fell 70% and that when NIH3T3 cells, among others, were serum starved, the levels of both proteins decreased by 30%. In both cases, ectopically expressed, GFP-tagged proteins were translocated from the nucleoli as first reported by Hanada *et al* (24).

These observations suggested that regulating the steady-state levels of the heterodimer could be one mechanism for regulating the levels of rDNA transcription. However, this model could only be true if rDNA transcription was dependent upon the heterodimer. Support of this model came from the observations of Hanada *et al*. that antibodies to PAF53 blocked specific rDNA transcription (24). However, as discussed above, neither of the yeast homologs of PAF53 or PAF49 were found to be essential for rDNA transcription. Additional evidence in support of this model came from CRISPR/Cas9 studies that demonstrated that both subunits were essential for mammalian cell proliferation (30) and our own that PAF53 was essential (51,53). However, these studies did not shed light upon the mechanism(s) that regulate the steady-state levels of the proteins.

In our study of the roles of PAF53 in rDNA transcription, we found that the heterodimerization domain was essential for activity. This suggested that the interaction with PAF49 might be essential for activity. Thus, when we initiated an investigation of the role(s) of PAF49 in rDNA transcription, we sought to determine if the knockdown of PAF49 affected the association of PAF53 with Pol I. In our first experiments to examine this question, we found that the auxin-induced knockdown of PAF49 resulted in the rapid decrease in total PAF53 levels, not just PAF53 associated with Pol I. Interestingly, we did not find that the auxin-induced degradation of PAF53 had a significant effect on PAF49 levels after 24 hours. It has been reported that the low molecular weight DNA intercalator, BMH-21, inhibits rDNA transcription and this induces the degradation of the high molecular weight subunit of Pol I (PolR1A) (54,55). As the knockout of PAF53 results in the inhibition of rDNA transcription, and the knockout of PAF49 caused the relatively rapid degradation of PAF53, we determined if the auxin-induced degradation of either subunit had an effect on the steady state-levels of several of the subunits of Pol I, including PolR1A. We did not find any evidence for a significant decrease in any of the other subunits of Pol I. We did find that knocking down PAF53 for 72hr. did cause a 70% decrease in the steady-state level of PAF49.

The auxin-induced degradation of PAF49 occurs through the activation of the TIR1-auxin dependent F-box protein and results in the ubiquitination and subsequent proteasome-dependent degradation of the targeted protein. As predicted, we found that the proteasome inhibitor MG-132 inhibited the auxin-dependent degradation of both AID-PAF53 and AID-PAF49. More importantly, when cells expressing AID-PAF49 were treated with MG-132 and IAA, the degradation of PAF53 was also inhibited. While not conclusive, this demonstrates that the degradation of PAF49 leads to the degradation of PAF53 and suggests that the steady-state levels of PAF53 in a cell are, to a very significant degree, regulated by the levels of its partner, PAF49.

We have carried out preliminary experiments to confirm that endogenous PAF53 was being targeted for degradation because of depletion of PAF49-AID, and not the result of an artifact of the auxin-Tir1 system. In these experiments, we used a pool of 3 siRNAs against PAF49 to knockdown PAF49. We found that knockdown of PAF49 via using siRNAs caused the attenuation of the steady-state levels of PAF53. These results support our model that levels of PAF49 play a significant role in regulating the steady-state levels of PAF53.

Based on the results from our previous studies and the experiments performed above, we hypothesize that PAF49 is required to allow PAF53 to bind to Pol I and function during transcription. Interestingly, the dimerization domain of yA49 is not required for the protein to function in transcription. On the other hand, the dimerization domain of its mammalian homolog, PAF53, is required for rDNA transcription and cell proliferation (51). If PAF53 could bind to Pol I independent of dimerization, then the dimerization domain would not be essential for transcription like its yeast homolog. Since the dimerization domain is required for function, PAF49 must be required for PAF53 to bind to Pol I and facilitate transcription. Further, we showed that after 3hrs of IAA treatment in PAF53-AID cells, the levels of PAF49 were not affected (Figure 4a). While reciprocal experiments in PAF49-AID

cells demonstrated a significant decrease in the levels of PAF53 (Figure 4b-c). We hypothesize that PAF49 is more stable in the absence of PAF53 because it can bind to Pol I in the absence of PAF53. In contrast, PAF53 gets quickly degraded because without PAF49, it cannot bind to the Polymerase and is vulnerable to degradation. Future studies will be performed to test the model that dimerization between PAF49 and PAF53 is required for PAF53 to bind to Pol I and function during transcription.

## Experimental Procedures

### Cell Culture, Transfection, Selection and Analysis

HEK293 cells (ATCC) were cultured as recommended in DMEM containing 10% FBS and Invitrogen Antibiotic-Antimycotic and the indicated inhibitors. Cells were routinely passed at 1:4 dilutions every third day and were not used after the 18th passage. Transfection of 60% confluent HEK293 cells was carried out as described (49,88) using PEI (117). After eight hours, the medium was changed. When selecting for stable transfections, the selection antibiotic was added 48 hr. post transfection. The established cell line expressing NLS-TIR1 was described previously, as was the cell line expressing both NLS-TIR1 and AID-tagged PAF53 (51). The AID tag was introduced into the 5’ coding region of the PAF49 gene using CRISPR/Cas9 and microhomology-mediated end-joining (MMEJ) repair (56). A manuscript characterizing these cell lines is in preparation. In order to induce auxin-dependent degradation, 500μM indole acetic acid (IAA) was added to the culture medium. IAA (Abcam) was made fresh by dissolving in water immediately before use.

### Western Blotting

Cell lysis and western blotting was carried out as described (126) using antibodies to PAF53 (48). PAF49, Rrn3, PolR1A and PolR1B and PolR1F (TWISTNB, RPA43). were either raised to recombinant proteins in our laboratory (26,57,58) or obtained from Proteintech Group and recognize both the human and mouse proteins. The antibodies from our laboratory have been validated previously (26,27,57). The depletion of endogenous PAF53 and PAF49 results in a loss of immunoreactivity for both proteins.

### RT-qPCR

RT-qPCR was used to determine the relative quantification of the mRNAs for PAF53 and PAF49. The results were analyzed by ΔΔCt, in which the expression or abundance ratio of the target gene mRNA in a sample was normalized to the abundance of PPPID mRNA, to allow for comparison between samples (59).

### Statistical Analysis

All experiments were reproduced at least three times with three technical replicates each time. Quantitative results that required comparisons between groups were subject to statistical analysis using two-tailed Student’s t test for two groups or one-way ANOVA followed by Dunnett’s multiple comparison test to determine significant differences among more than two groups. Data met assumptions of the tests (*i*.*e*., normal distribution, similar variance). Normality of our data was determined via a D’Agostino-Pearson omnibus normality test and a Shapiro-Wilk normality test. To determine whether the variance differed between groups, an F test was performed when comparing two groups and a Bartlett’s and Brown-Forsythe test was used to compare the variance of more than two groups

## Author Contributions

R. M., K. R., and L. I. R. conceptualization; R. M., K. R., and L. I.

R. investigation; R. M. and K.R. methodology; R. M. and L. I. R. writing-review and editing; L. I. R. funding acquisition; L. I. R. writing-original draft.

## Acknowledgements

This work was supported by National Institutes of Health Grants GM069841 (to L. I. R.), HL077814 (to L. I. R.), and F31CA250352 (to R.M.), OCAST Grant

HR15-166 (to L. I. R.), Presbyterian Health Foundation Grant 20201584 (to L.I.R.), and by the University of Oklahoma (to L.I.R.). *The authors declare that they have no conflicts of interest with the contents of this article*. The content is solely the responsibility of the authors and does not necessarily represent the official views of the National Institutes of Health.

## The abbreviations used are

Pol: polymerase
AID: auxin-inducible degron
IAA: indole acetic acid
ANOVA: analysis of variance
tWH: tandem-winged helix

## References Cited

1. Bateman, E., and Paule, M. R. (1986) Regulation of eukaryotic ribosomal RNA transcription by RNA polymerase modification. Cell 47, 445–450

2. Fath, S., Milkereit, P., Peyroche, G., Riva, M., Carles, C., and Tschochner, H. (2001) Differential roles of phosphorylation in the formation of transcriptional active RNA polymerase I. P Natl Acad Sci USA 98, 14334–14339

3. Peyroche, G., Milkereit, P., Bischler, N., Tschochner, H., Schultz, P., Sentenac, A., Carles, C., and Riva, M. (2000) The recruitment of RNA polymerase I on rDNA is mediated by the interaction of the A43 subunit with Rrn3. The EMBO journal 19, 5473–5482

4. Milkereit, P., and Tschochner, H. (1998) A specialized form of RNA polymerase I, essential for initiation and growth-dependent regulation of rRNA synthesis, is disrupted during transcription. The EMBO Journal 17, 3692–3703

5. Buttgereit, D., Pflugfelder, G., and Grummt, I. (1985) Growth-dependent regulation of rRNA synthesis is mediated by a transcription initiation factor (TIF-IA). Nucleic Acids Res 13, 8165–8180

6. Mahajan, P. B., Gokal, P. K., and Thompson, E. A. (1990) Hormonal regulation of transcription of rDNA. The role of TFIC in formation of initiation complexes. The Journal of biological chemistry 265, 16244–16247

7. Brun, R. P., Ryan, K., and Sollner-Webb, B. (1994) Factor C*, the specific initiation component of the mouse RNA polymerase I holoenzyme, is inactivated early in the transcription process. Molecular and cellular biology 14, 5010–5021

8. Schnapp, A., Schnapp, G., Erny, B., and Grummt, I. (1993) Function of the growth-regulated transcription initiation factor TIF-IA in initiation complex formation at the murine ribosomal gene promoter. Mol Cell Biol 13, 6723–6732

9. Moorefield, B., Greene, E. A., and Reeder, R. H. (2000) RNA polymerase I transcription factor Rrn3 is functionally conserved between yeast and human. P Natl Acad Sci USA 97, 4724–4729

10. Miller, G., Panov, K. I., Friedrich, J. K., Trinkle-Mulcahy, L., Lamond, A. I., and Zomerdijk, J. C. (2001) hRRN3 is essential in the SL1-mediated recruitment of RNA Polymerase I to rRNA gene promoters. EMBO J 20, 1373–1382

11. Yuan, X., Zhao, J., Zentgraf, H., Hoffmann-Rohrer, U., and Grummt, I. (2002) Multiple interactions between RNA polymerase I, TIF-IA and TAF(I) subunits regulate preinitiation complex assembly at the ribosomal gene promoter. EMBO Rep 3, 1082–1087

12. Cavanaugh, A. H., Hirschler-Laszkiewicz, I., Hu, Q., Dundr, M., Smink, T., Misteli, T., and Rothblum, L. I. (2002) Rrn3 phosphorylation is a regulatory checkpoint for ribosome biogenesis. J Biol Chem 277, 27423–27432

13. Cavanaugh, A. H., Evans, A., and Rothblum, L. I. (2008) Mammalian Rrn3 is required for the formation of a transcription competent preinitiation complex containing RNA polymerase I. Gene Expr 14, 131–147

14. Aprikian, P., Moorefield, B., and Reeder, R. H. (2001) New model for the yeast RNA polymerase I transcription cycle. Molecular and cellular biology 21, 4847–4855

15. Stepanchick, A., Zhi, H., Cavanaugh, A. H., Rothblum, K., Schneider, D. A., and Rothblum, L. I. (2013) DNA binding by the ribosomal DNA transcription factor rrn3 is essential for ribosomal DNA transcription. The Journal of biological chemistry 288, 9135–9144

16. Liljelund, P., Mariotte, S., Buhler, J. M., and Sentenac, A. (1992) Characterization and mutagenesis of the gene encoding the A49 subunit of RNA polymerase A in Saccharomyces cerevisiae. P Natl Acad Sci USA 89, 9302–9305

17. Nakagawa, K., Hisatake, K., Imazawa, Y., Ishiguro, A., Matsumoto, M., Pape, L., Ishihama, A., and Nogi, Y. (2003) The fission yeast RPA51 is a functional homolog of the budding yeast A49 subunit of RNA polymerase I and required for maximizing transcription of ribosomal DNA. Genes Genet Syst 78, 199–209

18. Gadal, O., Mariotte-Labarre, S., Chedin, S., Quemeneur, E., Carles, C., Sentenac, A., and Thuriaux, P. (1997) A34.5, a nonessential component of yeast RNA polymerase I, cooperates with subunit A14 and DNA topoisomerase I to produce a functional rRNA synthesis machine. Molecular and cellular biology 17, 1787–1795

19. Beckouet, F., Labarre-Mariotte, S., Albert, B., Imazawa, Y., Werner, M., Gadal, O., Nogi, Y., and Thuriaux, P. (2008) Two RNA polymerase I subunits control the binding and release of Rrn3 during transcription. Molecular and cellular biology 28, 1596–1605

20. Huet, J., Buhler, J. M., Sentenac, A., and Fromageot, P. (1975) Dissociation of two polypeptide chains from yeast RNA polymerase A. P Natl Acad Sci USA 72, 3034–3038

21. Engel, C., Plitzko, J., and Cramer, P. (2016) RNA polymerase I-Rrn3 complex at 4.8 A resolution. Nat Commun 7, 12129

22. Jennebach, S., Herzog, F., Aebersold, R., and Cramer, P. (2012) Crosslinking-MS analysis reveals RNA polymerase I domain architecture and basis of rRNA cleavage. Nucleic acids research 40, 5591–5601

23. Geiger, S. R., Lorenzen, K., Schreieck, A., Hanecker, P., Kostrewa, D., Heck, A. J., and Cramer, P. (2010) RNA polymerase I contains a TFIIF-related DNA-binding subcomplex. Molecular cell 39, 583–594

24. Hanada, K., Song, C. Z., Yamamoto, K., Yano, K., Maeda, Y., Yamaguchi, K., and Muramatsu, M. (1996) RNA polymerase I associated factor 53 binds to the nucleolar transcription factor UBF and functions in specific rDNA transcription. The EMBO journal 15, 2217–2226

25. Yamamoto, K., Yamamoto, M., Hanada, K., Nogi, Y., Matsuyama, T., and Muramatsu, M. (2004) Multiple protein-protein interactions by RNA polymerase I-associated factor PAF49 and role of PAF49 in rRNA transcription. Molecular and cellular biology 24, 6338–6349

26. Hannan, R. D., Hempel, W. M., Cavanaugh, A., Arino, T., Dimitrov, S. I., Moss, T., and Rothblum, L. (1998) Affinity purification of mammalian RNA polymerase I. Identification of an associated kinase. J Biol Chem 273, 1257–1267

27. Hannan, K. M., Rothblum, L. I., and Jefferson, L. S. (1998) Regulation of ribosomal DNA transcription by insulin. The American journal of physiology 275, C130–138

28. Penrod, Y., Rothblum, K., Cavanaugh, A., and Rothblum, L. I. (2014) Regulation of the association of the PAF53/PAF49 heterodimer with RNA polymerase I. Gene

29. Penrod, Y., Rothblum, K., Cavanaugh, A., and Rothblum, L. I. (2015) Regulation of the association of the PAF53/PAF49 heterodimer with RNA polymerase I. Gene 556, 61–67

30. Wang, T., Birsoy, K., Hughes, N. W., Krupczak, K. M., Post, Y., Wei, J. J., Lander, E. S., and Sabatini, D. M. (2015) Identification and characterization of essential genes in the human genome. Science 350, 1096–1101

31. Bertomeu, T., Coulombe-Huntington, J., Chatr-Aryamontri, A., Bourdages, K. G., Coyaud, E., Raught, B., Xia, Y., and Tyers, M. (2018) A High-Resolution Genome-Wide CRISPR/Cas9 Viability Screen Reveals Structural Features and Contextual Diversity of the Human Cell-Essential Proteome. Mol Cell Biol 38

32. Rothblum, L. I., Rothblum, K., and Chang, E. (2016) PAF53 is essential in mammalian cells: CRISPR/Cas9 fails to eliminate PAF53 expression. Gene

33. Chen, S., Seiler, J., Santiago-Reichelt, M., Felbel, K., Grummt, I., and Voit, R. (2013) Repression of RNA polymerase I upon stress is caused by inhibition of RNA-dependent deacetylation of PAF53 by SIRT7. Molecular cell 52, 303–313

34. Hannan, K. M., Kennedy, B. K., Cavanaugh, A. H., Hannan, R. D., Hirschler-Laszkiewicz, I., Jefferson, L. S., and Rothblum, L. I. (2000) RNA polymerase I transcription in confluent cells: Rb downregulates rDNA transcription during confluence-induced cell cycle arrest. Oncogene 19, 3487–3497

35. Zatsepina, O. V., Bouniol-Baly, C., Amirand, C., and Debey, P. (2000) Functional and molecular reorganization of the nucleolar apparatus in maturing mouse oocytes. Developmental biology 223, 354–370

36. Seither, P., Zatsepina, O., Hoffmann, M., and Grummt, I. (1997) Constitutive and strong association of PAF53 with RNA polymerase I. Chromosoma 106, 216–225

37. Blattner, C., Jennebach, S., Herzog, F., Mayer, A., Cheung, A. C., Witte, G., Lorenzen, K., Hopfner, K. P., Heck, A. J., Aebersold, R., and Cramer, P. (2011) Molecular basis of Rrn3-regulated RNA polymerase I initiation and cell growth. Genes & development 25, 2093–2105

38. Philippi, A., Steinbauer, R., Reiter, A., Fath, S., Leger-Silvestre, I., Milkereit, P., Griesenbeck, J., and Tschochner, H. (2010) TOR-dependent reduction in the expression level of Rrn3p lowers the activity of the yeast RNA Pol I machinery, but does not account for the strong inhibition of rRNA production. Nucleic acids research 38, 5315–5326

39. Schneider, D. A., and Nomura, M. (2004) RNA polymerase I remains intact without subunit exchange through multiple rounds of transcription in Saccharomyces cerevisiae. P Natl Acad Sci USA 101, 15112–15117

40. Bier, M., Fath, S., and Tschochner, H. (2004) The composition of the RNA polymerase I transcription machinery switches from initiation to elongation mode. FEBS letters 564, 41–46

41. Fath, S., Kobor, M. S., Philippi, A., Greenblatt, J., and Tschochner, H. (2004) Dephosphorylation of RNA polymerase I by Fcp1p is required for efficient rRNA synthesis. The Journal of biological chemistry 279, 25251–25259

42. Hirschler-Laszkiewicz, I., Cavanaugh, A. H., Mirza, A., Lun, M., Hu, Q., Smink, T., and Rothblum, L. I. (2003) Rrn3 becomes inactivated in the process of ribosomal DNA transcription. J Biol Chem 278, 18953–18959

43. Bodem, J., Dobreva, G., Hoffmann-Rohrer, U., Iben, S., Zentgraf, H., Delius, H., Vingron, M., and Grummt, I. (2000) TIF-IA, the factor mediating growth-dependent control of ribosomal RNA synthesis, is the mammalian homolog of yeast Rrn3p. EMBO Rep 1, 171–175

44. Fath, S., Milkereit, P., Podtelejnikov, A. V., Bischler, N., Schultz, P., Bier, M., Mann, M., and Tschochner, H. (2000) Association of yeast RNA polymerase I with a nucleolar substructure active in rRNA synthesis and processing. The Journal of cell biology 149, 575–590

45. Yamamoto, R. T., Nogi, Y., Dodd, J. A., and Nomura, M. (1996) RRN3 gene of Saccharomyces cerevisiae encodes an essential RNA polymerase I transcription factor which interacts with the polymerase independently of DNA template. The EMBO journal 15, 3964–3973

46. Schlosser, A., Bodem, J., Bossemeyer, D., Grummt, I., and Lehmann, W. D. (2002) Identification of protein phosphorylation sites by combination of elastase digestion, immobilized metal affinity chromatography, and quadrupole-time of flight tandem mass spectrometry. Proteomics 2, 911–918

47. Schnapp, A., Pfleiderer, C., Rosenbauer, H., and Grummt, I. (1990) A growth-dependent transcription initiation factor (TIF-IA) interacting with RNA polymerase I regulates mouse ribosomal RNA synthesis. EMBO J 9, 2857–2863

48. Dundr, M., Hoffmann-Rohrer, U., Hu, Q., Grummt, I., Rothblum, L. I., Phair, R. D., and Misteli, T. (2002) A kinetic framework for a mammalian RNA polymerase in vivo. Science 298, 1623–1626

49. Wang, T., Wei, J. J., Sabatini, D. M., and Lander, E. S. (2014) Genetic screens in human cells using the CRISPR-Cas9 system. Science 343, 80–84

50. Sadian, Y., Baudin, F., Tafur, L., Murciano, B., Wetzel, R., Weis, F., and Muller, C. W. (2019) Molecular insight into RNA polymerase I promoter recognition and promoter melting. Nat Commun 10, 5543

51. McNamar, R., Abu-Adas, Z., Rothblum, K., and Rothblum, L. I. (2019) Inducible degron-dependent depletion of the RNA polymerase I associated factor PAF53 demonstrates it is essential for cell growth and allows for the analysis of functional domains. J. Biol. Chem., In Press

52. Lee, D. H., and Goldberg, A. L. (1998) Proteasome inhibitors: valuable new tools for cell biologists. Trends Cell Biol 8, 397–403

53. Rothblum, L. I., Rothblum, K., and Chang, E. (2017) PAF53 is essential in mammalian cells: CRISPR/Cas9 fails to eliminate PAF53 expression. Gene 612, 55–60

54. Colis, L., Peltonen, K., Sirajuddin, P., Liu, H., Sanders, S., Ernst, G., Barrow, J. C., and Laiho, M. (2014) DNA intercalator BMH-21 inhibits RNA polymerase I independent of DNA damage response. Oncotarget 5, 4361–4369

55. Peltonen, K., Colis, L., Liu, H., Trivedi, R., Moubarek, M. S., Moore, H. M., Bai, B., Rudek, M. A., Bieberich, C. J., and Laiho, M. (2014) A targeting modality for destruction of RNA polymerase I that possesses anticancer activity. Cancer Cell 25, 77–90

56. Lin, D. W., Chung, B. P., Huang, J. W., Wang, X., Huang, L., and Kaiser, P. (2019) Microhomology-based CRISPR tagging tools for protein tracking, purification, and depletion. J Biol Chem 294, 10877–10885

57. Hannan, R. D., Cavanaugh, A., Hempel, W. M., Moss, T., and Rothblum, L. (1999) Identification of a mammalian RNA polymerase I holoenzyme containing components of the DNA repair/replication system. Nucleic acids research 27, 3720–3727

58. O’Mahony, D. J., Xie, W. Q., Smith, S. D., Singer, H. A., and Rothblum, L. I. (1992) Differential phosphorylation and localization of the transcription factor UBF in vivo in response to serum deprivation. In vitro dephosphorylation of UBF reduces its transactivation properties. J Biol Chem 267, 35–38

59. VanGuilder, H. D., Vrana, K. E., and Freeman, W. M. (2008) Twenty-five years of quantitative PCR for gene expression analysis. Biotechniques 44, 619–626

